# Cross-talk between active DNA demethylation, resetting of cellular metabolism and shoot apical growth in poplar bud break

**DOI:** 10.1101/122119

**Authors:** Daniel Conde, Mariano Perales, Anne-Laure Le Gac, Christopher Dervinis, Matias Kirst, Stéphane Maury, Pablo González-Melendi, Isabel Allona

## Abstract

Annual dormancy-growth cycle is a developmental and physiological process essential for the survival of temperate and boreal forests. Seasonal control of shoot growth in woody perennials requires specific genetic programs integrated with the environmental signals. The environmental-controlled mechanisms that regulate the shift between winter dormancy to growth promoting genetic program are still unknown. Here, we show that dynamics in genomic DNA methylation (gDNA) levels regulate dormancy-growth cycle in poplar. We proved that the reactivation of cell division in the apical shoot that lead bud break process in spring, is preceded by a progressive reduction of gDNA methylation in apex tissue. We also identified that the induction in apex tissue of a chilling-dependent poplar *DEMETER-LIKE 10* (*PtaDML10*) DNA demethylase precedes shoot growth reactivation. Transgenic poplars showing down-regulation of PtaDML8/10 caused delayed bud break. Genome wide transcriptome and methylome analysis and data mining revealed the gene targets of active DML-dependent DNA demethylation genetically associated to bud break. These data point to a chilling dependent-DEMETER-like DNA demethylase controlling the genetic shift from winter dormancy to a condition that promotes shoot apical vegetative growth in poplar.

## Introduction

Temperate and boreal perennials are distributed around the world in those regions in which four seasons are well defined based on their specific weather conditions along the year. Hence these tree species have developed complex molecular mechanism to adapt their annual cycle with the specific environmental conditions in each season, to guarantee their survival, sexual reproduction and seed set in adult trees. Winter dormancy is the strategy used by perennials in temperate and boreal regions to avoid cold damage. During winter dormancy, growth is inactivated in all meristems of the tree, i.e., the shoot apical meristem (SAM), the axiliary meristem, and the stem vascular cambium. Winter dormancy begins in autumn with an initial stage of ecodormancy, when the growth of the SAM and vascular cambium is arrested by short days (SD) and cold temperature. During this period, the tree forms autumn apical buds to protect the SAM and simultaneously develops moderate cold tolerance. The second stage is the endodormancy, when trees acquire maximum adaptation to the cold. These two states, ecodormancy and endodormancy, were described by (1). During this period, a chilling requirement needs to be fulfilled for growth to resume. Once the chilling period is fulfilled, the trees enter in the ecodormancy again, allowing bud break and the recovery of active growth under growth-promoting environmental conditions of spring. Annual cycle of temperate trees has been reviewed in (2).

Efforts are being focus on elucidating the winter dormancy control, since it determines several economically important traits in perennials, among them tree annual growth rate and fruit development. The main molecular players in dormancy regulation identified so far are: FLOWERING LOCUS T (FT) mediating the photoperiod signaling that leads to growth cessation at dormancy entry (2–4) and FD-LIKE1 (FDL1) and ABA INSENSITIVE 3 (ABI3) controlling bud maturation and cold acclimation in poplar (5). Maintenance of dormancy require the CENTRORADIALIS-LIKE1 (CENL1) regulator in poplar (6) and the regulation of the chromatin function by histone modifications associated to the function of DAM genes in peach (7, 8). At dormancy exit, EARLY BUD-BREAK1 (EBB1) is needed to promotes bud burst in poplar (9, 10).

Energy fluxes through the TCA cycle link the metabolic status of plants to the DNA methylation status (11). Cellular dynamics in NAD+/NADH ratio affects methylation status of plants (12). The reduction of cellular metabolism during the dormancy leads to a higher NAD+/NADH ratio that potentially would promote DNA methylation (11). According to this, in chestnut (*Castanea sativa*), the bud formation during winter dormancy and bud break during spring are accompanied by an increment and a decrease, respectively, of gDNA methylation (13). Similarly in apple (*Malus × domestica*) the transition from dormant bud to fruit set is accompanied by a progressive decrease of gDNA methylation (14). It has been proposed that the mechanisms of winter dormancy in trees and vernalization in herbaceous plants show similarities (8, 15). According to this, DNA hypomethylation resulted in early flowering or a reduced vernalization requirement to promote flowering in annual plants and to bolt tolerance in sugar beet (16–18).

DNA methylation refers to the addition of a methyl group in the fifth position of cytosine (5mC) and promotes changes in gene expression (19). DNA methylation occurs and is maintained by three DNA methyltransferases in plants that recognize different methylation contexts (CG, CHG and CHH, where H = A, C or T) (20). An important feature of DNA methylation is its reversibility, which can occur passively after de novo synthesis of unmethylated strands during DNA replication, or actively through the actions of enzymes that remove the methylated bases. In Arabidopsis there are four DEMETER-LIKE DNA demethylases (DMLs) (21). Active demethylation by DMLs enables the expression of genes in response to biotic and abiotic stresses in Arabidopsis (22, 23). Moreover, DMLs have been associated with developmental transitions during fruit ripening in tomato and nodule development in Medicago truncatula (24, 25).

Here, we addressed the role of genomic DNA methylation during dormancy-growth cycle regulation in poplar. We showed that the reactivation of shoot apical growth is preceded by reduced genomic DNA methylation in apex tissue while *DEMETER-LIKE DNA demethylase 10* (*PtaDML10*) is induced at the apex after a chilling fulfilment during ecodormancy. Our functional analysis showed that an active DML-dependent DNA demethylation is necessary for poplar bud break. Through genome wide transcriptome and methylome analysis and data mining, we identified the gene targets for the active DML-dependent DNA demethylation genetically associated to bud break. These data point to DEMETER-mediated DNA demethylation in controlling environmentally-induced developmental stage transition that shift from winter dormancy to a condition that promotes shoot apical vegetative growth.

## Results and Discussion

### Bud break is preceded by a reduced genomic DNA methylation in poplar apex while *PtaDML10* is induced during ecodormancy and bud break

To understand the temporal pattern of genomic DNA (gDNA) methylation during the transition from dormancy to growth, we profiled DNA methylation by high-resolution HPLC in poplar trees SAM growing under natural conditions, weekly from January to the time of bud break in April, (Fig. 1*A*). In this profile, a stage of gDNA hypermethylation in February was followed by a period of marked hypomethylation, with minimum levels detected on March 24. Subsequently, bud break (stages 1, 2 and 3 of apical bud development, Fig. 1 *A* and *B*), coincided with increased 5mC levels. Similar pattern of 5mC confer plasticity in environmentally-controlled responses such as vernalization in sugar beet (19). Thus, this particular gDNA methylation pattern is a common readout generated by the physiological state of the shoot apical cells during poplar dormancy and sugar beet vernalization.

**Figure 1.**
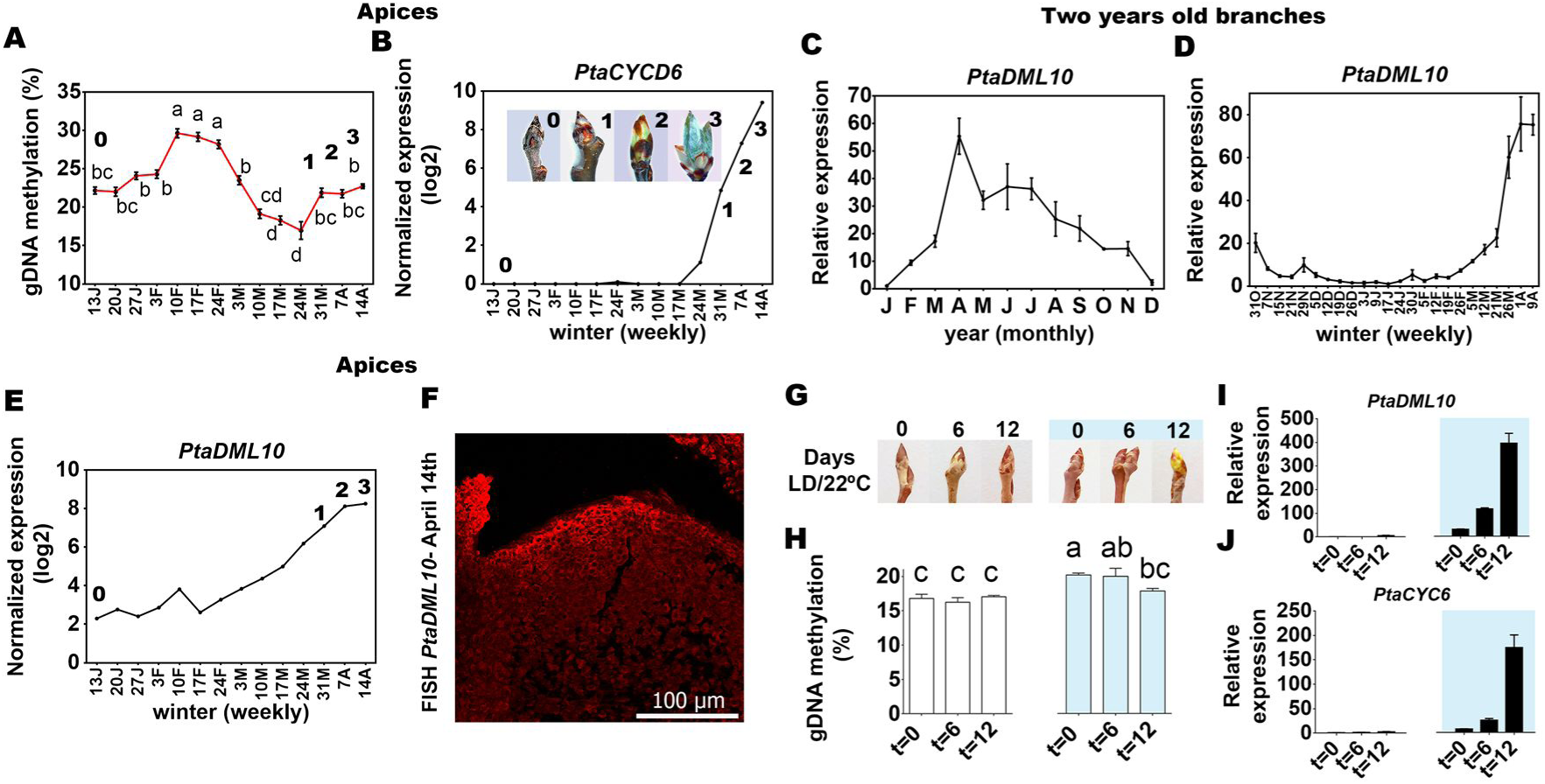
Analysis of global DNA methylation patterns and DNA demethylase mRNA expression from dormancy to bud break in poplar stem and apices. **(A)** DNA methylation levels (mean values +/− SE, n=4) in apical meristems quantified by HPLC in poplar growing under natural conditions from Jan 13 weekly to budburst on Apr 14. Each biological sample consists of a pool of n=28 dissected apices. Two independent hydrolyses were performed in each biological replicate and two HPLC analyses were carried out for each hydrolysis. Significant differences at each time point were analysed by ANOVA and TUKEY test. Points with different letters showed significant differences (P<0.01). Numbers indicate the different developmental stages of the apical bud during the period of this experiment according to the bud burst stages defined by UPOV (1981). **(B)** Normalized expression of *PtaCYCD6* obtained by RNA sequencing in poplar apices growing under natural conditions from Jan 13 to bud break on Apr 14. Figures and numbers shown inside the *PtaCYCD6* profile indicate the developmental stage of apices as described in (A). **(C,D)** Quantitative RT-PCR analysis of DML10 in 2-year-old poplar branches: (C) over the year and (D) during winter dormancy from Oct 31 weekly to bud break on Apr 9. Plotted values and error bars are fold-change means ± SD recorded in three technical replicates. **(E)** Normalized expression of *PtaDML10* was determined by RNA sequencing in poplar apices growing under natural conditions from Jan 13 to bud break on Apr 14. Numbers shown inside the *PtaDML10* profile indicate the developmental stage of apices as described in (A). **(F)** Fluorescence immunodetection of *PtaDML10* in 20 μm WT apical bud sections collected at bud break on Apr 14 from poplar trees growing under natural conditions. **(G)** Phenological observation of dormant apices of cuttings taken from poplar trees growing under natural conditions before and after fulfilling the chilling requirement and transferred to LD and 22°C conditions. After 12 days in conditions of LD, 22°C, most postchilling cuttings started bud break, while the apices of prechilling cuttings remained at stage 0. Prechilling cuttings are indicated in white and postchilling cuttings in blue. **(H)** DNA methylation levels (mean values +/− SE, n=8) in dormant apical meristems quantified by HPLC in dormant apices of pre- and postchilling cuttings grown under natural conditions and after 6 and 12 days of exposure to LD, 22°C. Two independent hydrolyses were performed for each biological replicate and two HPLC analyses were carried out for each hydrolysis. Significant differences at each time point were analysed by ANOVA and TUKEY test. Points with different letters showed significant differences (P<0.05). **(I,J)** Quantitative RT-PCR analysis of *PtaDML10* and *PtaCYCD6* in dormant apices of pre- and postchilling cuttings grown under the same conditions as in (G). Plotted values and error bars are fold-change means ± SD recorded in three technical replicates. Two biological samples were analysed at each point consisting of a pool of n=15 dissected apices.

To elucidate whether the decrease in gDNA methylation precedes or follows cell cycle reactivation, we examined the expression of D-type cyclin *PtaCYCD6:*1, a marker of growth reactivation in poplar apical meristem (26). *PtaCYCD6:*1 mRNA levels rose from March 17 until the last time point April 14 (bud stage 3, based on the scale established in (27, 28) (Fig. 1*B*) indicating that most likely this decrease in gDNA methylation level is produced by an active DNA demethylation mechanism, because it occurs previously to the reactivation of cell cycle that lead bud break.

This scenario led us to identify poplar DNA demethylases, through phylogenetic and protein sequence analysis. We identified three putative *DMLs* genes in poplar, *PtaDML6, PtaDML8* and *PtaDML10*, containing all domains and amino acids conserved in DMLs, essential for active 5mC excision (Fig. S1). To elucidate if one of these poplar DMLs could be involved in the reduction of gDNA methylation that precedes bud break, we analysed their annual expression pattern (monthly) in stems, and their expression pattern during winter dormancy (weekly) in stems and apices. Annual expression patterns of the three genes in poplar stems revealed that only *PtaDML8* and *PtaDML10* showed highest levels during vegetative growth, with PtaDML8 expression peaking in June during the highest growth period and PtaDML10 expression peaking at the time of bud break (April) (Fig. 1*C* and Fig. S2*A*). In the expression profile for poplar stems and apices, *PtaDML10* induction was observed in late winter dormancy (end of February) with maximum expression levels produced at bud break (April 1 and 9), while *PtaDML8* expression remained constant in stems and apices (Fig. 1 *D* and *E* and Fig. S2 *B* and *C*). These observations are consistent with previous transcriptome analyses indicating the increased *PtaDML10* expression during dormancy release, the transition from endo- to ecodormancy that precedes the bud break, in poplar (29).

Additionally, *PtaDML10* mRNA was localized by FISH in poplar apices. The fluorescence signal was detected in the L1 layer and central SAM, and also in the tips of leaf primordia at bud break (April 14) (Fig. 1*F*). This spatio-temporal localization is similar to that reported for EARLY BUD-BREAK 1 (EBB1), a putative APETALA2/ethylene responsive transcription factor in poplar (30). EBB1 overexpression causes early bud break in poplar, and activates CYCD3 genes in Japanese pear, whereas its down-regulation delays this process in poplar (30, 31). Relatively higher expression of EBB1 in L1 indicates the importance of this layer in the reactivation of SAM growth in spring.

In order to dissect whether the activation of *PtaDML10* and the reduction of gDNA methylation occur during endo- or ecodormancy stages, cuttings were taken from trees grown in their natural environment in both conditions, a set of cuttings taken from trees that had not fulfilled the chilling requirement (endodormant cuttings) and a set of cuttings from trees that did fulfil the chilling requirement (ecodormant cuttings). Both sets were transferred to growth promoting-conditions, long day (LD) at 22°C, in a growth chamber and phenological observations were made at 0, 6 and 12 days after transfer (Fig. 1*G*). 85% of the ecodormant apical buds on day 12 reached stages 1 or 2 (27, 28), indicating the initiation of bud break (Fig. 1*G*). The gDNA methylation level in the ecodormant apical buds was higher than in the prechilling ones, but decreased significantly 12 days after transfer to growth-promoting environment, when bud break process began (Fig. 1*H*). However, the gDNA methylation levels in endodormant apical buds did not show changes during the same period (Fig. 1*H*). Accordingly, *PtaDML10* RNA levels remained very low in endodormant apices, while it expression was gradually induced in ecodormant apices, reaching the maximum expression when the bud break began (after 12 days under growth-promoting environment) (Fig. 1*I*). The resumption of growth in the ecodormant samples after 12 days was indicated by the expression of *PtaCYCD6* (Fig. 1*J*). In summary, the results of this experiment showed: in endodormant apical buds there is not initiation of bud break after 12 days under growth promoting-conditions, there are neither changes in gDNA methylation nor induction of *PtaDML10* during this period. In contrast, ecodormant apical buds initiated bud break after 12 days under growth-promoting environment, and this process is accompanied by both, a significant reduction of gDNA methylation level and a huge induction of PtaDML10 expression. All together indicate that dormancy release is necessary for PtaDML10 induction during the process of bud break, and suggest that it could be implicated in the gDNA demethylation that precedes bud break.

Collectively, the expression pattern suggests that *PtaDML8* would play a role during the vegetative growth period, particularly in stem tissues, and that *PtaDML10* could be implicated in the gDNA methylation reduction observed in apical buds previous to the reactivation of cell division that leads to the bud break process. Induction of *PtaDML10* occurs four weeks earlier than *PtaCYCD6*:1, together with the higher induction of *PtaDML10* mRNA in L1 at this time, suggesting its role in the reestablishment of growth after winter dormancy.

### Active DML-dependent DNA demethylation is required for bud break

Functional studies were performed by generating *PtaDML8/10* knock-down plants (KD) (Fig. S3). RT-PCR analyses showed that both *DML* genes were downregulated in the transgenic plants due to the high similarity existing between *PtaDML8* and *PtaDML10*. We examined phenological variables in two lines showing low-level *PtaDMLs* expression, KD2 and KD5. When grown under LD and 22°C conditions, in vitro or glasshouse cultured transgenic plants were phenotypically similar to WT plants. Moreover, these lines showed no differences compared to WT in cessation of growth elongation and bud formation when transferred from LD to short day (SD) conditions (Fig. 2 *A* and *B*). These observations rule out a role of these DMLs in dormancy entry. WT and KD plants were subjected to 0, 4 and 8 week of SD conditions at 4°C and bud burst was monitored in LD conditions at 22°C using the scale shown in Fig. 2*C* (27, 28). Without chilling, t=0, there was no bud break, indicating that these two DMLs do not affect dormancy release. More significant differences in bud burst were observed after 4 weeks (Fig. 2*D*). Thus, lower levels of *DMLs* expression led to a significant delay in growth reactivation in the transgenic trees after dormancy, pointing to a role of these *DMLs* in bud break phenology. In addition, through HPLC of genomic DNA from dissected apices, 5mC levels 4 and 2% higher were detected in KD2 and KD5, respectively than in WT (Fig. 2*E*).

**Figure 2.**
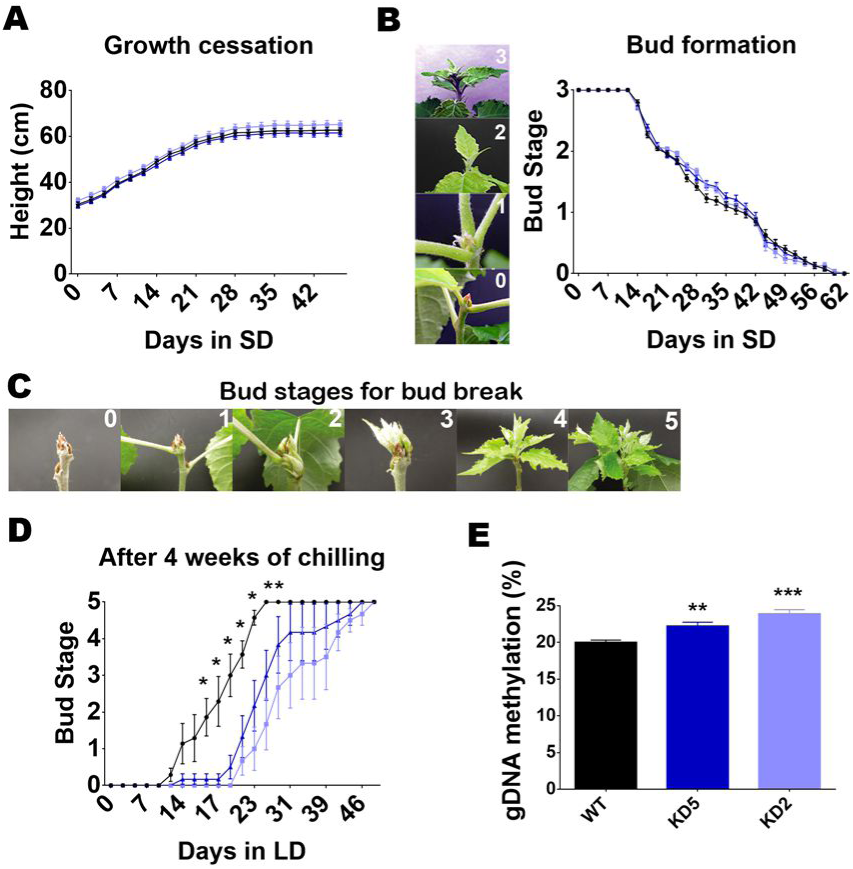
An active DML-dependent DNA demethylation is necessary for bud break. **(A,B)** Growth cessation and bud set in response to SDs, 19°C. WT, KD2 and KD5 plants were grown in conditions of LD16h, 22°C for 4 weeks and then height and bud scores were monitored in response to SDs, 19°C. Mean heights (cm) ± SE and mean bud scores are represented for the wild type (n = 24), KD2 (n = 23) and KD5 (n = 25). **(C)** Apex stages during bud break. The scale used to monitor growth reactivation after inducing apical bud formation and fulfilling the chilling requirement was developed by (22). **(D)** Timing of bud burst after dormancy. Timing of bud burst was examined in response to LD and warm temperature after dormancy induction. WT, KD2 and KD5 plants were treated as shown in (A,B). Following a shift to SD and 4°C, plants were transfer to LD and 22°C at three different times: t=0, t=4wk and t=8wk. Here we show apical growth reactivation monitored in the 4wk experiment. Mean bud burst scores ± SE were measured for the wild type (n = 7), KD2 (n = 7) and KD5 (n = 7). Significant differences between KDs and WT were analysed using the Wilcoxon Rank Sum test in R: * P<0.05. **(E)** DNA methylation level in apices of WT, KD2 and KD5. DNA methylation levels (mean values +/− SE, n=4) in dormant apical meristems quantified by HPLC in WT, KD2 and KD5. Shoot apical meristems were collected at the time point close to bud burst. Each biological sample consists of a pool of n=15 dissected apices. Two independent hydrolyses were performed for each biological replicate and two HPLC analyses for each hydrolysis. Significant differences between KDs and WT were assessed by the Student’s t-test: ** P<0.01; *** P<0.001.

This loss of function studies confirmed that active 5mC demethylation is necessary for bud break. Even though *PtaDML8/10* knock-down plants have reduced expression of *PtaDML10* and *PtaDML8*, the earlier described spatio-temporal expression patterns suggest that PtaDML10 mainly participates in the chilling-dependent genomic DNA demethylation needed for successful bud break after winter dormancy.

### Target genes of the active DML-dependent DNA demethylation during bud break

The delayed bud break phenology observed in our *PtaDML8/10* KD lines led us to hypothesize that *DMLs* downregulation causes inefficient reactivation of growth promoting genes or/and, that these lines were unable to turn off genes required for dormancy maintenance. To identify genes demethylated by PtaDMLs in poplar apices, WT, KD2 and KD5 lines were subjected to whole genome bisulfite sequencing and genome wide RNA sequencing transcriptome analysis. These data and the experimental flow through are provided in Fig. 3 and Dataset S1-3.

**Figure 3.**
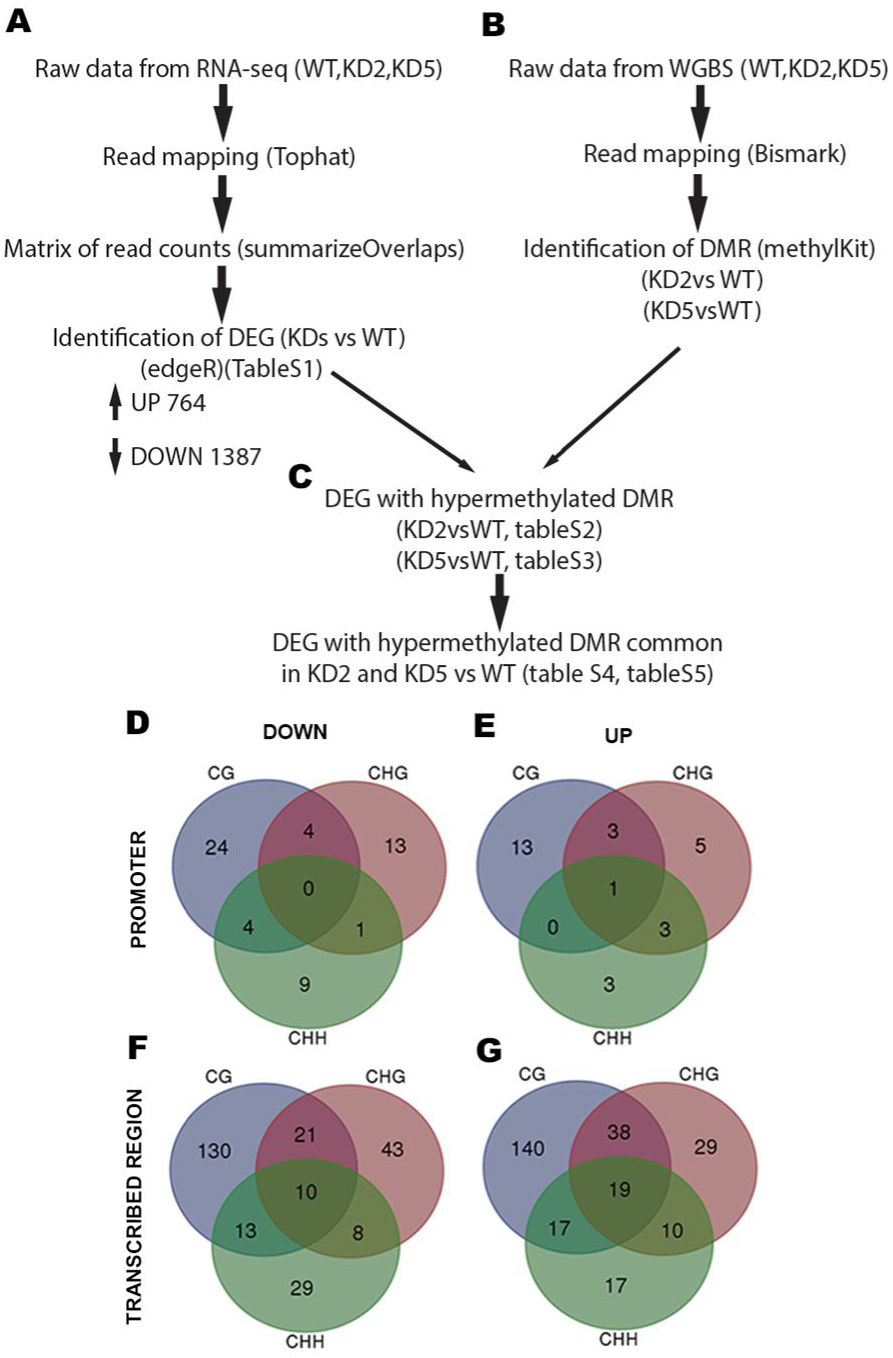
Analysis pipeline and identification of DEGs with hypermethylated DMR in the promoter or transcribed region in KD lines versus WT. **(A)** Flowchart of RNA-seq analysis pipeline. Transcriptome comparisons revealed 2151 differentially expressed genes (DEG) in KD lines compared to WT, among which 764 were upregulated and 1387 downregulated. **(B)** Flowchart of WGBS analysis pipeline. **(C)** Flowchart of the identification of DEGs with hypermethylated DMR in KDs vs. WT. The identification of DEGs with hypermethylated DMR either in their promoter or transcribed region was performed in KD2 vs. WT and KD5 vs. WT. Next, common DEGs in both datasets were identified. **(D, E)** Methylation contexts in DEGs with hypermethylated DMR in the promoter are shown in a Venn diagram. A set of 83 common DEGs, 55 down- (D) and 28 up-regulated (E), with hypermethylated DMR in the promoter region were detected mostly in CG and CHG contexts. **(F, G)** Methylation contexts in DEGs with hypermethylated DMR in the transcribed region are shown in a Venn diagram. We found 524 DEGs, 254 down- (F) and 270 up-regulated (G), with hypermethylated DMR common to both transgenic lines located mainly in a CG context.

We expected that the growth-promoting gene targets of the active DML-dependent DNA demethylation would be highly expressed in young proliferating aerial tissues and induced during bud break. Thus, among the DEGs that were hypermethylated in the three contexts either in their promoter or transcribed regions, and downregulated in KD plants compared to WT, we selected those that were relatively more expressed in young leaves and SAM. Among them, we pulled out the genes induced at the time of bud break based on our weekly RNA sequencing profiles in poplar apices growing under natural conditions from Jan 13 to bud break on Apr 14 available in geneAtlas (https://phytozome.jgi.doe.gov/pz/portal.html) (Dataset S4 and S5). As shown in Fig. 4 *A-C*, of the 19 and 60 genes identified respectively in the two sets, almost 75% grouped in the categories of four gene ontology biological processes related to cell metabolism: protein translation, chloroplast metabolism and light reaction, secretory pathway and mitochondrial metabolism. An open question for further studies will be to explore the role in bud break of the 7 genes ascribed here to chromatin remodelling and transcription regulation biological process. Remarkably, among these genes we found poplar homologues to Arabidopsis genes with a pivotal role in meristem development and maintenance, like *SHOOTMERISTEMLESS* (32), *ULTRAPETALA* 1 (33) or *KNOTTED1-like homeobox gene 6* (34), indicating the active DML-dependent DNA demethylation promotes reactivation of meristem activity.

**Figure 4.**
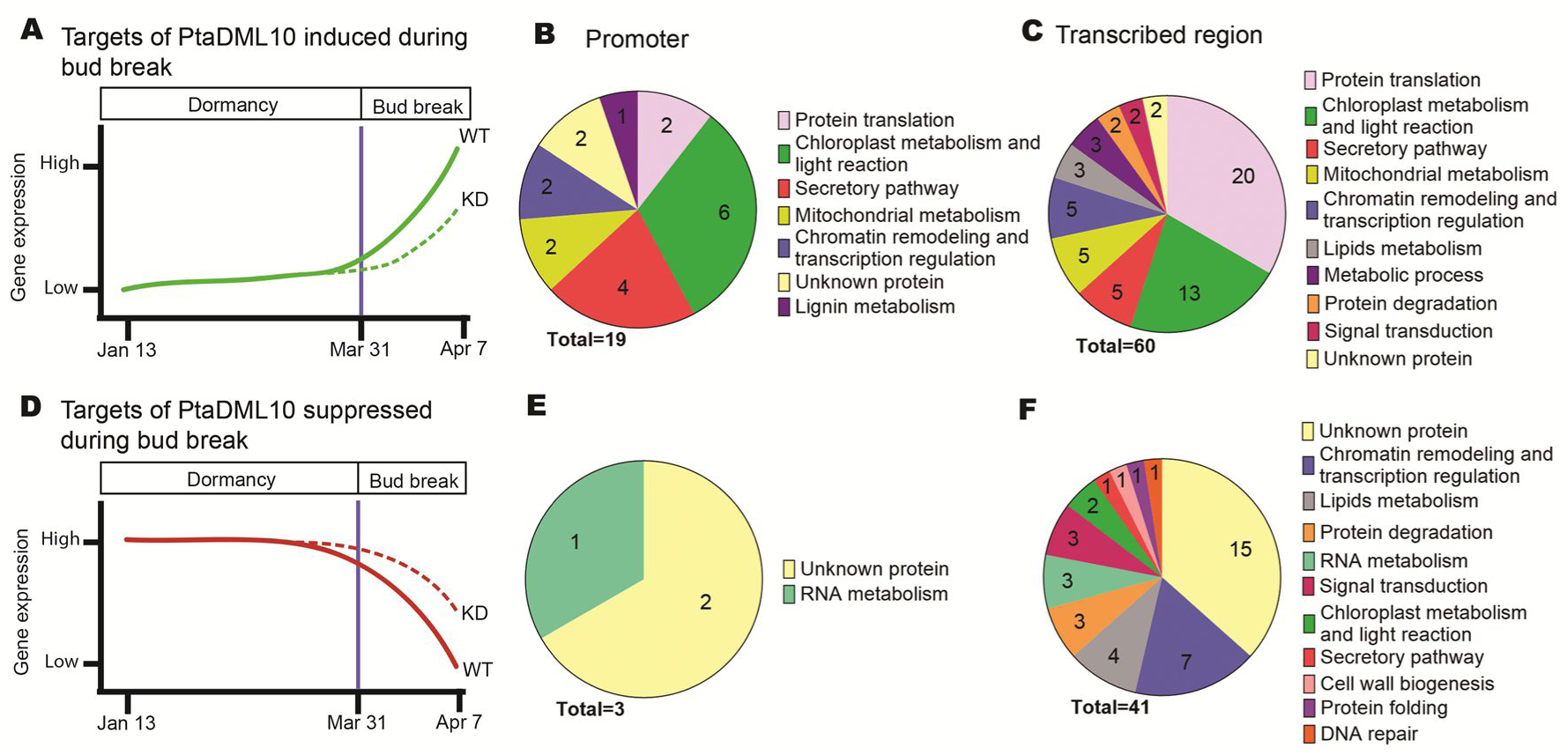
Gene targets of active DML-dependent DNA demethylation during bud break. **(A)** Graphical model showing the inefficient reactivation of growth promoting genes in KD plants during bud break. **(B,C)** Biological processes for DEG genes hypermethylated in their promoter region **(B)** or in their transcribed region **(C)** and downregulated in KD lines compared to WT, primarily expressed in young leaves or SAM and induced during bud break. **(D)** Graphical model showing the inefficient repression of dormancy genes in KD plants during bud break. **(E,F)** Biological processes of DEG genes hypermethylated in their promoter region **(E)** or in their transcribed region **(F)**, upregulated in KD lines compared to WT and repressed during bud break.

To identify the genes required for dormancy maintenance repressed by active DML-dependent DNA demethylation, we selected from the sets of DEGs that were hypermethylated in the promoter or transcribed region, and upregulated in KD plants compared to WT, those that were suppressed during bud break in our RNAseq time series deposited in geneAtlas (Dataset S4 and S5). As shown in Fig. 4 *D-F*, of the 3 and 41 genes identified, respectively, more than 38% were unknown genes, indicating a lack of detailed knowledge about the regulation and maintenance of the dormant state. This may only be studied in perennial plants.

### Target genes of the active DML-dependent DNA demethylation and bud break phenology are genetically associated in poplar

A genome wide single nucleotide polymorphisms (SNPs) has been associated with phenotypic variation in several traits, including bud break, in a collection of 544 *Populus trichocarpa* individuals spanning much of natural latitudinal range of this specie (35). To identify common candidates controlling bud break, we merged our list of the active DML-dependent DNA demethylation targets and the list of genes showing SNPs that correlates with bud flush phenotypic variation in Evans et al. work (35). We found 15 genes associated with bud break, 9 of them were identified as growth-promoting gene targets of the active DML-dependent DNA demethylation showing a marked activation during the process of bud break in our temporal RNAseq dataset, with the exception of *KNAT6* which mRNA expression pattern remained unaltered during the winter (Fig. 5 A-H, Dataset S6). Among them, we identified 6 genes participating in pathways controlling protein homeostasis of the plant cell: One poplar homolog to Arabidopsis *CLP PROTEASE 5 (CLPP5)* that encodes a subunit of chloroplast protease complex ClpPR involved in chloroplast biogenesis and thylakoid protein homeostasis (36). One poplar homolog to Arabidopsis and Maize gene that encodes a plastidial RidA (Reactive Intermediate Deaminase A) required to the hydrolysis of the reactive and toxic intermediates generated from the serine and threonine metabolism in the chloroplast (37). Moreover, the disruption of RidA impaired photosynthesis and growth in Arabidopsis (37). One poplar homolog to Arabidopsis *TRANSLOCASE OF THE INNER MITOCHONDRIAL MEMBRANE 13 (TIM13)*, a member of the protein import apparatus needed for protein mitochondria homeostasis (38). Additionally, we found two poplar homologs to Arabidopsis genes encoding ribosomal subunits S8e and L19e required for protein translation (39) and one poplar homolog to Arabidopsis gene that encodes a member of the ubiquitin-dependent protein catabolic process (40). Jointly, these data indicate that the active DML-dependent DNA demethylation could promote plant cell growth and bud break by the activation of key regulators of the cellular protein homeostasis. In addition to this, we identified 3 target genes, genetically associated to bud break, that participate in shoot meristem growth control: One poplar homolog to Arabidopsis *NON-PHOTOTROPIC HYPOCOTYL 3 (NPH3)*, a blue light signaling transducer essential for shoot phototropism, leaf flattening and positioning in Arabidopsis (41-43). One poplar homolog to *PROTODERMAL FACTOR 1 (PDF1)*, a L1 layer secreted protein suggested to play a role in L1 cell specification (44). One poplar homolog to *KNOTTED1-like homeobox gene* 6 (*KNAT6*) required for shoot meristem activity and organ boundary formation (27). Accordingly, these data show that the active DML-dependent DNA demethylation drives bud break by activating blue light and L1 specific cell signalling, and overall meristem activity.

**Figure 5.**
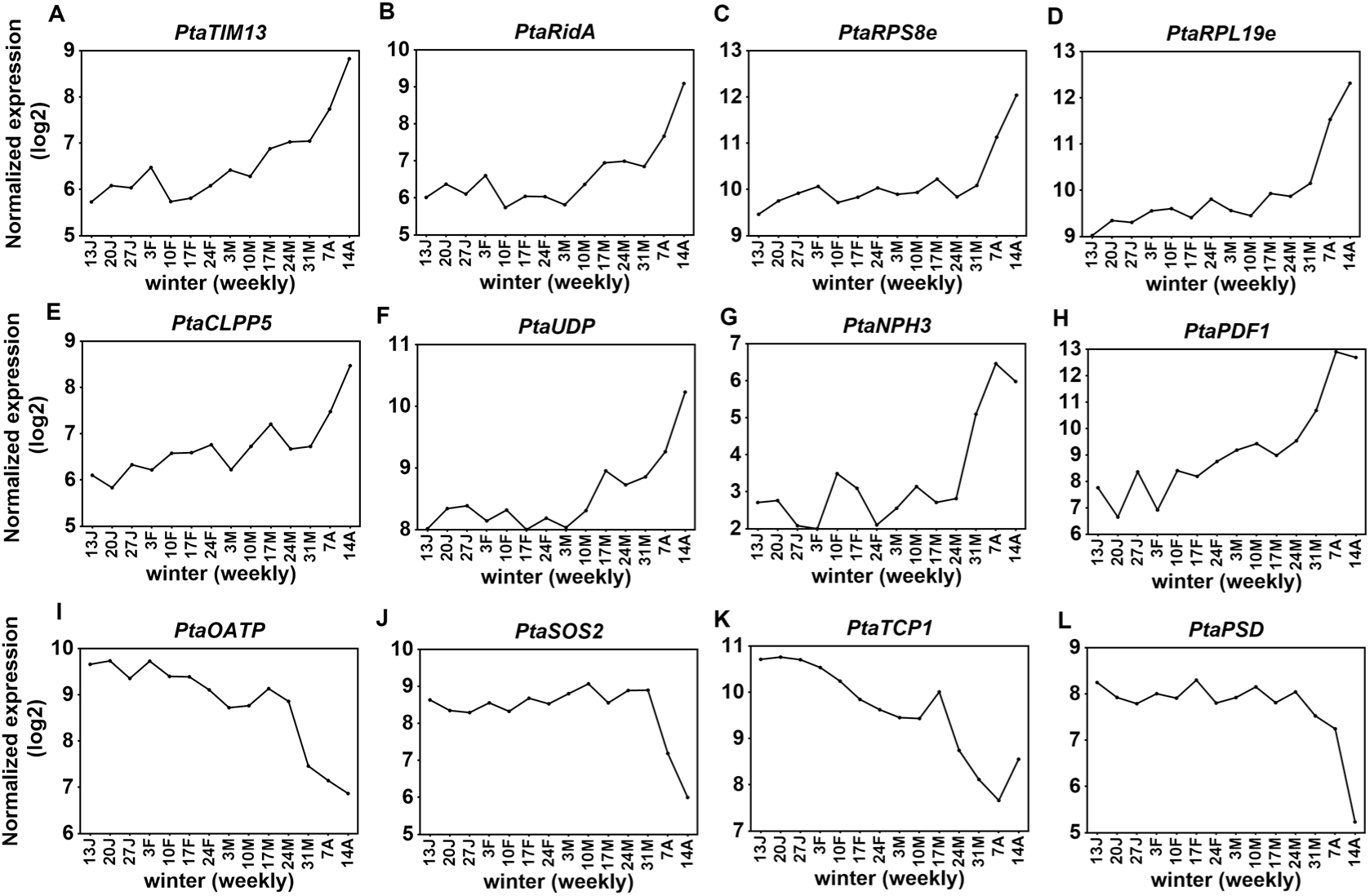
Normalized expression pattern of DMLs target genes genetically associated to bud break. **(A)** *TRANSLOCASE OF THE INNER MITOCHONDRIAL MEMBRANE 13 (PtaTIM13)*, **(B)** Reactive Intermediate Deaminase A (*PtaRIDA*), **(C,D)** Ribosomal S8e and L19e family genes (*PtaRPS8e* and *PtaRPL19e*), **(E)** *CLP PROTEASE 5 (PtaCLPP5)*, **(F)** A gene that encodes a ubiquitin-dependent protein catabolic process (*PtaUDP*), **(G)** *NON-PHOTOTROPIC HYPOCOTYL 3 (PtaNPH3)*, **(H)** *PROTODERMAL FACTOR 1 (PtaPDF1)*, **(I)** *ORGANIC ANION TRASPORTING POLIPEPTIDE (OATP)*, **(J)** *SALT OVERLY SENSITIVE 2 (SOS2)*, **(K)** *TWO PORE CHANNEL 1 (TCP1)*, **(L)** *PAUSED (PSD)*.

Genetically associated with bud break, we also identified 6 downregulated target genes, showing a decreasing pattern of mRNA accumulation in our temporal RNAseq dataset (Fig.5 IK, Dataset S6). Two of them having unknown function in Arabidopsis and Poplar. Within the remaining 4 genes, it is remarkable the identification of three poplar homolog to Arabidopsis and human proteins involved in regulation of ion homeostasis. SALT OVERLY SENSITIVE 2 (SOS2) a serine-threonin protein kinase that regulate sodium ion homeostasis (45), TWO PORE CHANNEL 1 (TCP1) a nonselective cation channel on the vacuole membrane (46) and the homolog to the ORGANIC ANION TRASPORTING POLIPEPTIDE (OATP) from humans (47)Moreover, a poplar homolog to the Arabidopsis PAUSED (PSD) importin β-family of nucleocytoplasmic transport receptors involved in exporting from the nucleus to the cytoplasm tRNAs, and others RNAs and proteins (48). Therefore, the active DML-dependent DNA demethylation could play a role in shifting from a winter osmotic stress to a regular growth promoting condition, and moreover readjusting the nuclear-cytoplasm trafficking needed to promote bud break.

In apple, Kumar et al. (2016) (14) found that reduced genomic DNA methylation during bud break affects the expression of genes involved in many cellular metabolic processes. Taken together these observations suggest a conserved mechanism controlling growth-dormancy cycles in deciduous trees. Although a role for *DEMETER-LIKE* genes in Arabidopsis embryo development has been long established (49), mutants in Arabidopsis DMLs show little or no ulterior developmental alterations (50,51,52), suggesting that active DNA demethylation is not critical for normal development in this species. The participation of these genes in postembryonic development has recently emerged in crops. Thus, it has been shown that active 5mC demethylation by DEMETER proteins is required for transcriptional reprogramming during tomato fruit maturation (30). Further, symbiotic organogenesis uses DEMETER-dependent epigenetic regulation to reprogram gene expression in specific host cells in *Medicago truncatula* (23), and, as observed here, this mechanism restores sensitivity to growth-promoting stimuli for bud break in poplar. To date, the molecular regulation of bud break is poorly understood. So far, only a molecular cascade triggered by EBB1 has been implicated in the regulation of this process (10, 30). However, no overlap was found when we crossed the active DML-dependent DNA demethylation direct target genes of with known direct targets of EBB1 (30). Hence, it seems that DEMETER and EBB1 mechanisms act via separate pathways to control bud break.

## Conclusion

Our data shows that a active DNA demethylation driven mainly by a chilling-dependent *PtaDML10* activity is necessary for bud break and occurs as two simultaneous processes: 1) by enabling the reactivation of key genes controlling protein homeostasis checkpoints, along with the activation of blue light and L1 specific cell signalling, and overall meristem activity; and 2) a process whereby those genes that are more highly expressed during bud dormancy are downregulated, including the resetting cellular osmotic conditions and nuclear-cytoplasm export. Collectively, these results show that an active DML-dependent DNA demethylation could act as molecular switch shifting from winter dormancy to a condition that promotes shoot apical vegetative growth.

## Materials and Methods

A detailed description of the materials and methods used in the paper are provided in SI Materials and Methods. The following items are included: phylogenetic analyses to identify poplar DML proteins; plant material and growth conditions; RT-PCR expression analysis; probe synthesis and fluorescence in situ hybridization (FISH); generating binary vector constructs and plant transformation; RNA sequencing in shoot apices; DNA methylation analysis and methylome analysis through whole genome bisulfite sequencing (WGBS).

## Acknowledgments

This study was supported by grants AGL2011-22625, AGL2014-53352-R, KBBE PIM2010PKB-00702, and COST Action FP0905 awarded to I.A.; M.P. and D.C. were supported by the Ramon y Cajal program of MINECO (RYC-2012-10194) and PCIG13-GA-2013-631630; M.K. holds a US NSF PGRP IOS-1444543 grant. Two short stays by D.C. in the lab of S.M. in Orléans, France were funded by grants from the Consejo Social of the Universidad Politécnica de Madrid and Universidad Politécnica de Madrid. The authors also thank A. Delaunay (University of Orléans, France) for technical advice and assistance with genomic DNA methylation determination by HPLC, Phil Wigge for his helpful comments and suggestions during this work and C. Grunau and C. Chaparro (IHPE, Perpignan, France) from the RTP3E (http://rtp-3e.wix.com/rt3e) for the use of their local bioinformatics cluster.

## SI Materials and Methods

### Phylogenetic analyses to identify poplar DML proteins

Arabidopsis DML protein sequences were retrieved through the TAIR10 website (www.arabidopsis.org). These sequences were used as queries to BLAST the Populus trichocarpa proteome (Phytozome v11, https://phytozome.jgi.doe.gov/pz/portal.html) and identify homologue proteins in this organism. These protein were aligned using MAFFT (E-INS-i algorithm) (1) and edited in Jalview 2.8 (2). The best evolutionary model of the alignment was highlighted using ProtTest 2.4 (3). The maximum likelihood tree was inferred using the RAxML tool (4) with 100 bootstrap replicates and visualized at the iTOL website http://itol.embl.de (5). Bootstrap values were generated with 100 replicates.

The conserved regions of Arabidopsis and poplar DMLs were assembled at the PROSITE website (http://prosite.expasy.org/mydomains/).

### Plant material and growth conditions

To explore 5mC profiles from winter dormancy to bud break, 28 apical buds were collected weekly from 7 hybrid poplar trees (*Populus tremula* × alba INRA clone 717 1B4) at the Centre for Plant Biotechnology and Genomics (CBGP) in Pozuelo de Alarcón, Madrid (3°49’W, 40°24’N) over the period January 13 to April 14, 2015. Each week, genomic DNA was extracted from the 28 dissected apices for HPLC.

Annual expression patterns for poplar *PaDML8* and *PaDML10* were determined in 2-year-old poplar branches (*Populus alba*) growing under natural conditions in Madrid (Spain). The expression profiles of *PtaDML8* and *PtaDML10* throughout winter were measured in 2-year-old poplar branches and apical buds of hybrid poplar (*Populus tremula × alba* INRA clone 717 1B4) at CBGP.

To examine PtaDML10 induction in pre-chilling and post-chilling conditions, cuttings were collected from five 6-year-old hybrid poplars growing under natural conditions. In a first experiment, cuttings were harvested on December 11, when the buds were in a dormant state and the chilling requirement had not been fulfilled (endodormant cuttings). In a second experiment, cuttings were collected on March 7, when the chilling requirement had been fulfilled but buds were still in an ecodormant state. Thirty-six cuttings (approximately 12 cm long) per tree were collected for each experiment. 120 cuttings were placed vertically in empty blue pipette tip racks filled with water to cover the bottom 5 cm of the cuttings. Each box contained 15 cuttings (three cuttings of each selected tree). Cuttings in racks were transferred to the growth chamber under LD conditions (16 h light/) at 22°C. Apical buds were scored as six developmental stages of bud burst (stage 0 to stage 5) according to (6, 7).

For phenological assays, plantlets of *in vitro*-cultivated wild type and two *PtaDML8-10* RNAi lines, KD2 and KD5, were transferred to 3.5 L pots containing blond peat, pH 4.5. Before any photoperiod treatment, the plants were grown in a chamber under controlled LD conditions (16h light, 21°C, 65% relative humidity and 150 μmol m^-2^ s^-1^ light intensity). After 4 weeks, the plants were transferred to SD conditions (8h light, 19°C, 65% relative humidity and 150 μmol m^-2^ s^-1^ light intensity) for 12 weeks to induce apical bud formation. Bud formation progression was scored as stage 3 (fully grown apex) to 0 (fully formed apical bud) according to (8). Dormant plants were kept under the conditions SD, 100 μmol m^-2^ s^-1^ light intensity and 4°C for 0, 4 and 8 weeks. Monitoring of bud burst after dormancy induction and chilling was conducted in LD conditions at 22°C. Regrowth was scored for the six developmental stages of bud burst (stage 0 to stage 5) according to (6, 7). Significant differences in apex state were identified through pairwise comparisons between KD and WT plants using the Wilcoxon Rank Sum test after Shapiro-Wilk confirmation of the non-normal distribution of data. All statistical tests were run in R (www.r-project.org).

We repeated the phenological experiment to induce winter dormancy induction and release (with 4 weeks of chilling requirement) in WT, KD2 and KD5 plants. When growth resumed, the topmost (apical) bud of each plant was collected when the first visual changes in bud development were evident. Buds with morphological changes indicating initial stages of bud break were removed, so all apical buds were collected at stage 0. We collected 30 apical buds from 30 plants for each of the WT, KD2 and KD5 genotypes; 15 were used for RNA extraction and sequencing, and 15 for genomic DNA extraction for HPLC and whole genome bisulfite sequencing analyses.

### RT-PCR expression analysis

Total RNA extraction, single-stranded cDNA synthesis, primer design, real-time PCRs and data analysis were performed as described previously (9). The gene-specific primers used are described in Fig S4A.

### Probe synthesis and fluorescence in situ hybridization (*FISH*)

Apical buds for this experiment were harvested on April 14, 2015. A 271 base pair (bp) fragment of the 5’ portion of the untranslated terminal region (UTR) of *PtaDML10* DNA was cloned into pGem-T (Promega, Madison, WI, USA). The probe was *in vitro* transcribed to obtain digoxigenin-UTP (DIG) labelled RNA probes using T7 and Sp6 RNA polymerases for the sense and the anti-sense mRNA probes, respectively (Roche Applied Science, Mannheim, Germany).

Apices were fixed and preserved as previously described in (10) before embedding in paraffin wax. The specimens were washed twice in PBS (30 min each) and dehydrated in a series of 30%, 50%, 70% (1 h each) and 95% ethanol overnight. Specimens were then washed in 100% ethanol for 12 h. Ethanol was then replaced with a mixture of 50% histoclear (HC)-50% ethanol overnight and 100% HC for 12 h (twice). Vials containing the apices were filled up to a quarter of their volume with paraplast and incubated at 42°C for 12 h. Next, another quarter volume was added, followed by incubation at 60°C overnight. Paraplast in HC was replaced with melted paraffin wax (60°C), and this was replenished every 12 h for 5 days. Finally, the apices were placed in moulds and after solidification at room temperature they were kept at 4°C. Sections of 15-20 μm were obtained in a microtome Leica 2055 (Leica Microsystems, Wetzlar, Germany) and the paraffin removed in HC for 15 min and then in 100% ethanol for 15 min. Finally, the sections were rehydrated in a series of 96% and 70% ethanol for 10 min each and, finally, in PBS for 15 min.

After deparaffination, dehydration and rehydration, as previously described, bud sections were treated with 2% cellulase and proteinase K (1 μg ml^-1^) at 37°C, for 1 hour each. Next, 30 ng μl^-1^ of either the sense or the antisense probe diluted in the hybridization mixture (50% formamide(v/v), 10% dextrane sulphate (w/v), 10 mM Pipes, 1 mM EDTA, 300 mM NaCl, 1000 ng μl -1 tRNA) were applied to the sections and these incubated at 50°C overnight. After hybridization, the slides were washed in 1X saline sodium citrate buffer (SSC) for 30 min at 50°C. DIG detection and fluorescence visualization were carried out with a primary anti-DIG (Sigma) antibody diluted 1/3000 in 3% BSA in PBS for 1 h at room temperature, and a secondary anti-mouse 546 (ALEXA, Molecular Probes) antibody, applied 1/25 in 3% BSA in PBS for 1 h at room temperature in the dark. Images were acquired using a Leica TCS-SP8 confocal microscope under the laser excitation line of 561 nm. Gain and offset conditions were maintained in all captures for comparison. Images were acquired using a Leica TCS-SP8 confocal microscope.

### Generating binary vector constructs and plant transformation

A 419 bp fragment of Poplar *DML10* was amplified from *Populus alba* winter stem cDNA using gene specific primers with attB sites (Fig S4B). We followed the protocol described in (11) for RNAi fragment insertion in the hybrid poplar genome. Plantlets were generated in kanamycin-containing selection medium and independently transformed individuals (lines) were screened by reverse transcription (RT)-PCR to down-regulate *PtaDME10* expression.

### RNA sequencing in shoot apices

Expression profiles were examined by RNA sequencing (RNA-Seq) in dormant apical bud at the same time point, just before growth resumption after winter dormancy induction and release in *PtaDML10 KD2, PtaDML10 KD5* and wild-type hybrid poplars. Before RNA extraction, apices were dissected under a stereomicroscope to remove the leaflets surrounding the shoot apical meristem. RNA extraction was performed according to an earlier published protocol (9). Libraries were prepared using 1 μg of RNA in the NEBNext^®^ Ultra™ Directional RNA Library Prep Kit following the supplier’s instructions (New England Biolabs, Ipswich, MA, USA). Sequencing was done with an Illumina NextSeq500 in high-throughput mode (2×150 cycles) at the Interdisciplinary Center for Biotechnology Research at the University of Florida (Gainesville, FL, USA).

Pre-processing and assembly of sequence data were conducted using the web-based platform Galaxy at https://usegalaxy.org/ (12–14), which includes tools to remove reads containing N blurs, adapter sequences, and sequences shorter than 15 nucleotides. Quality control of reads was carried out with the FastQC Galaxy tool. Removal of rRNA, if needed, was performed locally with the SortMeRNA tool. The TopHat Galaxy tool was used to map clean reads to the *Populus tremula x P. alba* INRA clone 717-1B4 genome using the default settings. We used the *summarizeOverlaps* function (packages Genomic Alignments of R) to generate the matrix of read count for each gene from BAM files. Differentially expressed genes (DEG) lists were obtained using the function exactTest (edgeR package of R). For DEG analyses, *PtaDML10 KD2* and *PtaDML10 KD5* transcriptomes were considered as biological replicates when compared to WT, using a *FDR* < 0.05. Using this strategy, we tried to identify only genes that are consistently differentially regulated between *PtaDML10 KD* transgenic lines, detecting the biological impact of the transgene in down-stream regulation. Raw data for the RNAseq were deposited in Gene Expression Omnibus at NCBI under GEO accession number GSE87445.

### DNA methylation analysis

Genomic DNA methylation levels were quantified by HPLC in poplar apices dissected under a stereomicroscope to remove the leaflets surrounding the shoot apical meristem. Genomic DNA was purified from poplar samples and prepared for HPLC using a published method (15, 16) via RNase A digestion, phenol/chloroform extraction, and ethanol precipitation. Purified DNA was hydrolyzed into nucleosides using DNase I (700 U, Roche Diagnostics, Meylan, France), phosphodiesterase I (0.05 U, SerLabo Technologies, Entraigues sur la Sorgue, France), and alkaline phosphatase type III (0.5 U, Sigma-Aldrich, Saint-Quentin Fallavier, France) successively. Percentage genomic methylation was determined by HPLC using a GeminiTM column (150 9 4.6 mm, 5 lm, Phenomenex, Le Pecq, France) and isocratic mobile phase composed of 0.5 % methanol (v/v) and 5 mM acetic acid in water according to the methods of (16). Controls for this procedure were co-migration with commercial standards (Sigma-Aldrich), confirmation by enzyme restriction analysis (15), and tests for RNA contamination through HPLC detection of ribonucleosides. The methylcytosine percentage was calculated following the previously described method (17).

### Methylome analysis through whole genome bisulfite sequencing (*WGBS*)

gDNA extracted from apices of WT, KD2 and KD5 specimens just before bud break was sent to the Fasteris DNA sequencing service (https://www.fasteris.com) for sequencing after bisulfite treatment using the Hiseq 2500 platform. Unmethylated lambda DNA was added to the total DNA before bisulfite treatment to calculate the conversion rate, which was estimated as 94.16, 93.90 and 90.91 for WT, KD2 and KD5, respectively. The Galaxy platform of the University of Perpignan’s server was used for quality control of reads and to obtain the read alignment against the *Populus tremula x alba* INRA clone 717-1B4 genome using Bismark mapper (18). Before mapping, the last nucleotide of all reads was trimmed because of its lower sequencing quality (<28). In total, 19.02%, 19.04% and 18.36% of reads were mapped against the 717-1B4 genome in WT, KD2 and KD5, respectively. To identify differentially methylated regions (DMR) in KD2 vs. WT and KD5 vs. WT, the methylkit (19) package of R was used with a minimum difference of 15% in methylation level (R studio software, University of Perpignan’s server). We calculated the methylation status for cytosines with a minimum sequence depth of 10. A total of 14616377 (10.24%), 16559883 (11.61%) and 15097166 (10.58%) cytosines (% of cytosines in the genome) were evaluable in WT, KD2 and KD5, respectively. Raw data for whole genome bisulfite sequencing were deposited in Gene Expression Omnibus at NCBI, under GEO accession number GSE87445.

To identify which DEG with hypermethylated DMR in their promoter or transcribed regions showed relatively higher expression in young leaves and SAM, we first crossed them with genes identified in previously published transcriptome analysis performed in the roots, young and mature leaves, nodes and internodes of *Populus trichocarpa* reference genotype Nisqually-1. The expression patterns of the genes in the different tissues from this report were examined in the public database www.popgenie.org, and those mainly expressed in young leaves were separated. We also identified genes with the Arabidopsis homologue primarily expressed in SAM by crossing DEG with the expression browser tool selecting AtGenExpress_Plus-Extended Tissue Series at Arabidopsis eFP Browser (http://www.bar.utoronto.ca/). To identify which of those genes are induced or suppressed during bud break, we used expression data generated by RNA-seq in the experiment conducted in poplar apices from January 13 to April 14. These data are available at the phytozome11 website (https://phytozome.jgi.doe.gov/pz/portal.html).

## Supplemental information

**Figure S1.**
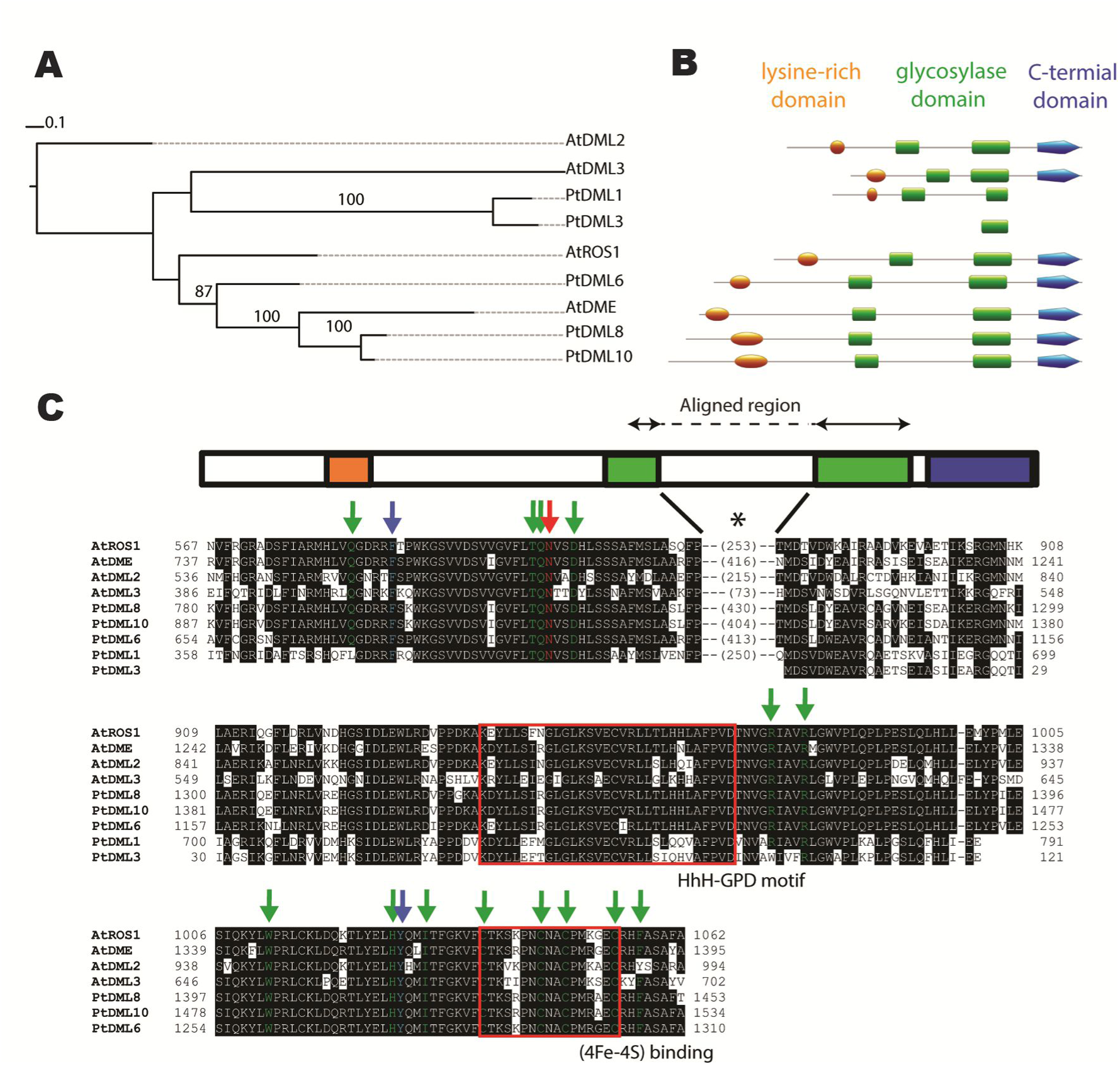
Phylogenetic tree analyses of DML proteins from Arabidopsis and poplar. (A) Maximum likelihood tree obtained using the RAxML tool with 100 bootstrap replicates. (Only bootstraps higher than 70 are shown). Arabidopsis protein sequences of DMLs were retrieved using the TAIR10 website (www.arabidopsis.org). These sequences were used as queries to BLAST the Populus trichocarpa proteome (Phytozome v10.3, www.phytozome.net) and identify homolog proteins in this organism. These proteins were aligned using MAFFT (E-INS-i algorithm) and edited in Jalview 2.8. The best evolutionary model of the alignment was highlighted using ProtTest 2.4. The maximum likelihood tree was inferred with the RAxML tool (Stamatakis, 2006) and visualized in the iTOL website http://itol.embl.de. Bootstrap values were generated with 100 replicates. **(B)** Scheme showing 5mC DNA glycosylase conserved domains in Arabidopsis and poplar. Domains were assembled in the PROSITE website (http://prosite.expasy.org/mydomains/). PtDML6, PtDML8 and PtDML10 conserve all the domains required for 5mC DNA glycosylase activity. Orange oval represents the lysine-rich domain. Green rectangles represent the motifs of the DNA glycosylase domain. Blue pentagon represents the c-terminal domain. **(C)** MAFF alignment of the 5mC DNA glycosylase domain of Arabidopsis and poplar. Black boxes indicate identical aminoacids. Black asterisk indicates the non-conserved linker sequence between the two motifs of the DNA glycosylase domain. Arrows indicate the position of the aminoacids necessary for the 5mC demethylation activity of AtROS1, AtDME and AtDML2-3 (green: required for base 5mC excision, blue: required for preference for 5mC over T mismatches, red: involved in control of context preference for 5mC). Red boxes indicate HhH-GPD and [4Fe–4S] motifs. PtaDML6, PtaDML8 and PtaDML10 show all the aminoacids that have been described to be essential for DMLs activity.

**Figure S2.**
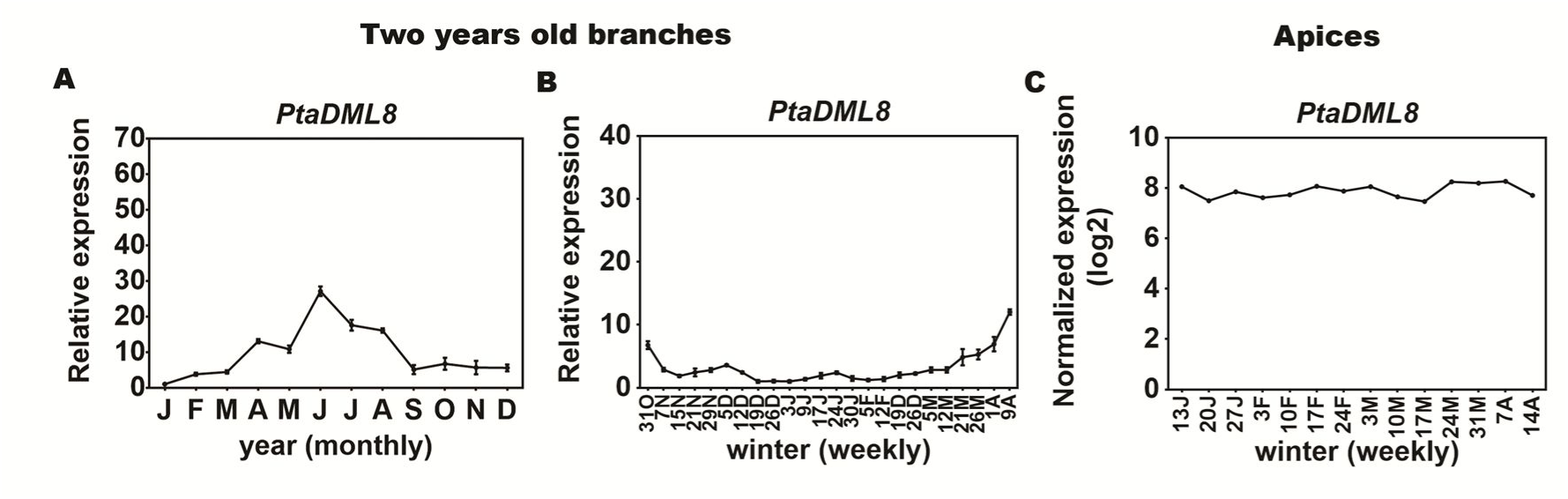
PtaDML8 mRNA expression from dormancy to bud break in poplar. **(A,B)** Quantitative RT-PCR analysis of DML8 in poplar 2-year-old branches: (A) along the year and (B) during winter dormancy from Oct 31 weekly to bud break on Apr 9. Plotted values and error bar are fold-change means ± SD recorded in three technical replicates. **(C)** Normalized expression of PtaDML8 was determined by RNA sequencing in poplar apices growing under natural conditions from Jan 13 to bud break on Apr 14.

**Figure S3.**
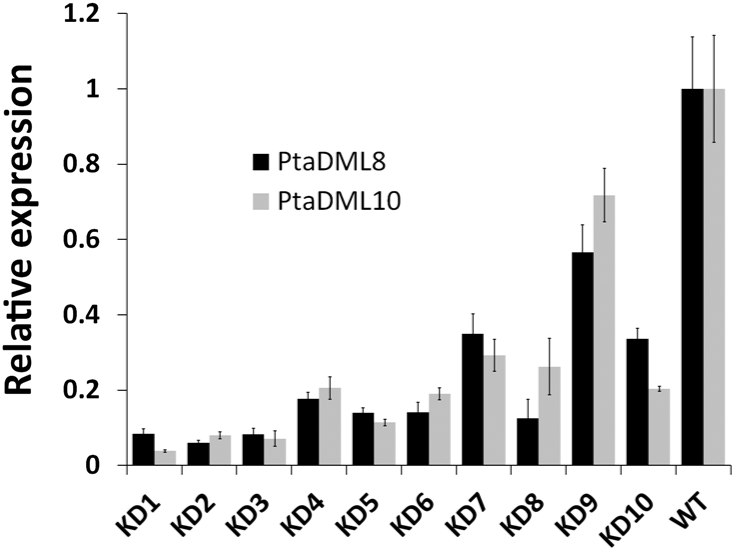
Characterization of PtaDML10 RNAi transgenic lines. Quantitative RT-PCR analysis of PtaDML8 and PtaDML10 in WT plantlet leaves. Samples were collected from plantlets growth in a greenhouse for 5 weeks in conditions of LD, 22°C. Plotted values and error bars are the fold-change means ± SD recorded in three technical replicates.

**Figure S4.**
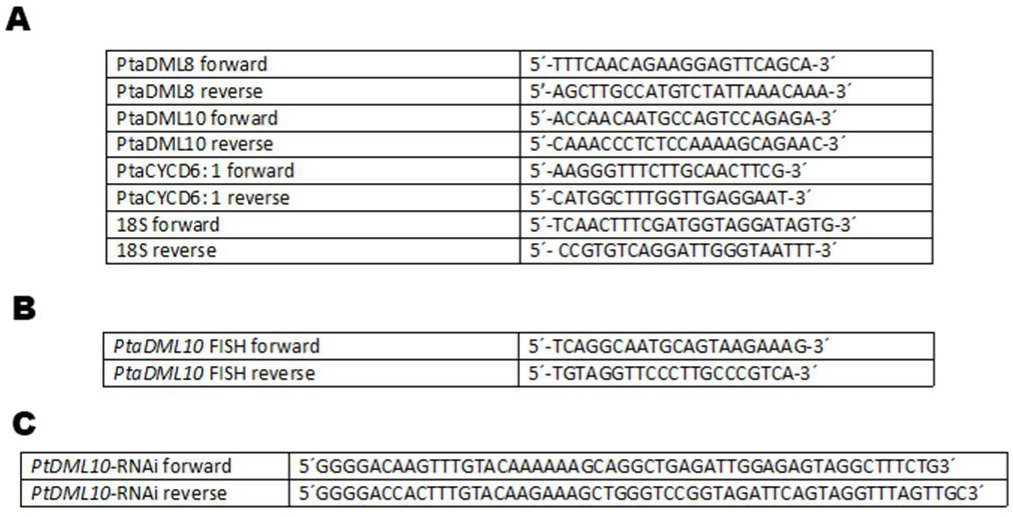
**(A)** The gene-specific primers used for the analyses of their expression by real-time PCR. **(B)** Primers used for amplify the DNA fragment used to generate RNA probe for *PtaDML10* mRNA FISH **(C)** Primers with attB sites used to generate the RNAi fragment for downregulated PtaDML10 gene expression.

